# Development of the Highly Sensitive Horse questionnaire to assess horses’ sensory processing sensitivity

**DOI:** 10.64898/2026.10.29.762739

**Authors:** Mélissa Peignier, Maya Bräm, Jenni Elise Kähkönen, Sabrina Briefer Freymond

## Abstract

It is well established that individuals vary in how they *respond* to situations, but might they also *perceive* their environment differently? Sensory Processing Sensitivity (SPS) is a multidimensional trait describing individual variation in the perception and response to sensory, emotional, and social stimuli. While extensively studied in humans, its presence in animals remains underexplored. We investigated whether horses might exhibit a comparable trait and developed the Highly Sensitive Horse questionnaire to assess equine SPS. A 104-item questionnaire was developed and refined using data from an online survey of 121 horses. The remaining 45 items were evaluated in an independent survey of 561 horses. Exploratory and confirmatory factor analyses identified a robust two-factor, 12-item structure reflecting dimensions similar to the human SPS dimensions of: ‘Overstimulation’, describing a tendency to become overwhelmed by intense or prolonged stimuli, and ‘Sensitivity to subtleties’, describing heightened perception of subtle environmental changes. The scale showed high internal consistency, excellent intra-rater and moderate inter-rater reliability. Although eSPS was associated with established behavioural traits, including fearfulness, it also explained unique behavioural variation beyond these traits. These findings suggest that horses differ in how they perceive their environment, providing a foundation to better understand equine behaviour, and opening new avenues for comparative sensory-processing research.

## Introduction

Individual animals and humans often differ in their behavioural responses to the same situation. When such between-individual variation is coupled with within-individual consistency across time and/or contexts, it is referred to as personality [1,2]. Numerous studies have demonstrated the role of personality in shaping fitness-related outcomes, including reproduction and survival [3–6]. However, most studies to date have focused on how individuals *respond* differently to similar situations, assuming that all subjects *perceive* the situation in the same way. In reality, perception itself is shaped by each organism’s unique sensory abilities and individual sensitivity [7].

In humans, Aron and Aron [8] identified a repeatable behavioural trait describing differences in how individuals perceive, process and react to the sensory, emotional or social stimuli of their environment. This trait, termed *Sensory Processing Sensitivity* (SPS), has since been conceptualised as a multidimensional personality construct [9–11]. Despite extensive research, the exact dimensional structure of SPS remains under debate. One framework describes SPS through four subdimensions summarized by the acronym DOES [12,13]: (i) D for *Depth of processing*, reflecting how deeply individuals process sensory information; (ii) O for *Overstimulation*, reflecting susceptibility to sensory overload; (iii) E for *Emotional reactivity*, reflecting the intensity of emotional responses; and (iv) S for *Sensitivity to subtleties*, reflecting awareness of subtle environmental details. Regardless of the precise dimensional structure, it is well established that SPS constitutes a behavioural trait rather than a pathology, disease, or mere enhancement of sensory acuity [14,15], with approximately 20 to 30% of the human population scoring high on the SPS trait [14]. While Sensory Processing Sensitivity has been extensively studied in humans, and animal personality research has expanded in recent decades, SPS in non-human animals remains largely unexplored.

Understanding how animals perceive and respond to their environment is critical to better predict their reactions. To date, only a couple of studies have investigated SPS in a non-human species. Bräm et al. [16,17] identified a canine SPS trait analogous to human SPS spanning overstimulation, emotional reactivity, and sensitivity to subtleties. The authors further demonstrated that dogs with high SPS scores are more prone to behavioural issues, particularly when (i) the owner uses aversive modes of communication, (ii) the owner has a lower SPS score than the dog, or (iii) negative punishment methods are used [17]. As dogs are an integral part of the daily life of many households, poor knowledge of their personality can compromise their well-being and the safety of humans and other animals around them. A similar concern arises for horses, which are increasingly considered companion animals, despite their historical classification as livestock [18–20]. Adapting human practices to the horse’s personality may help reduce accidents, improve their well-being and ultimately strengthen the human-animal relationship [21].

One previous study provided preliminary evidence for individual variation in equine sensitivity to sensory stimuli. Lansade et al. [22] presented horses with different stimuli in order to trigger the five different senses and reported that horses’ behavioural responses to sensory input were stable over time and across contexts. Yet, assessing all potential dimensions of SPS (i.e., sensory sensitivity, emotional reactivity, overstimulation, and depth of processing) through standardized behavioural tests is unrealistic for most private horse owners. To achieve a more comprehensive understanding of horse personality, questionnaires represent a useful complementary tool, despite being subject to potential bias [23]. Several horse personality questionnaires already exist and have been used successfully by horse owners [24–26]. Some of these questionnaires even contain items that potentially overlap conceptually with SPS. For example, ‘nervousness’ described by [27] as being ‘nervous about insects, noises, etc.’, may reflect differential sensitivity to external stimuli. In order to advance equine personality research, it is therefore essential to examine whether an equine SPS trait (eSPS) exists and how it relates to previously studied dimensions.

In the present study, we developed the Highly Sensitive Horse questionnaire to investigate the existence of an equine Sensory Processing Sensitivity trait. We selected items via semi-directed interviews with horse owners, and by using items from the validated Highly Sensitive Person, Highly Sensitive Child and Highly Sensitive Dog questionnaires [8,9,16] that could be applicable to horses. We subsequently reduced the number of items, validated them and analysed their latent structure through two international online surveys. We hypothesized that an eSPS-related behavioural trait, analogous to SPS in humans and dogs, could be assessed in horses using an owner-reported questionnaire, and that this trait would be at least partially independent from previously studied horse behavioural traits as has been observed in humans. The present study focused on the development and initial validation of the scale. Given the breadth of the analyses presented, further assessments of construct validity (e.g., standardized behavioural tests designed to assess eSPS) and investigation of links with other variables (e.g., breed, sex, detention mode) were beyond the scope of this paper.

## Results

### 1. Refinement of the questionnaire

The initial questionnaire we created consisted of 104 items and three selective questions, and was submitted online to horse owners, who could only answer for one horse (N = 121). We added the score of the three selective questions and assigned the top 20% of individuals (N = 24) to the potentially ‘highly sensitive horse’ group and the low 30% of individuals (N = 36) to the potentially ‘not highly sensitive horse’ group. These values were chosen based on the assumption that approximately 20% of humans exhibit high sensitivity and 30% exhibit low sensitivity [14].

Out of the 104 initial items, 15 were removed because more than 10% of respondents selected either ‘I don’t know’ or ‘Not applicable’ (Supplementary Table S1), thus suggesting that these items were not suitable or relevant for the broader equine population. Subsequently, 43 items were removed because they did not show a credible difference between the two groups in the Bayesian linear mixed-effect model (p < 0.09; Supplementary Table S2). Lastly, an examination of the correlation matrix between the 46 remaining items indicated two highly correlated items (r > 0.8). The item with the highest proportion of ‘I don’t know’ or ‘Not applicable’ answers was removed. In total, 45 items were retained. A complete list of retained and excluded items is provided in Supplementary Table S1.

### 2. Validation of the questionnaire

The 45-item questionnaire was submitted online again, together with demographic questions (e.g., sex, age, country of residence, type of training methods used, behavioural problems) and previously validated questions about other behavioural traits in horses [24]. A total of 481 participants completed the questionnaire, providing ratings for 561 horses (range: 1–7 horses per participant). Participants had a mean age of 44 years (range: 9–81 years), while horses had a mean age of 14 years (range: 1–31 years). Of the 481 participants, 243 identified as female, 237 as male, and one as non-binary. The sample comprised 259 French-speaking, 209 German-speaking, and 13 English-speaking participants. Among the horses, 277 were geldings, 267 were mares, and 17 were stallions.

First, exploratory factor analysis was performed on half of the sample to refine the items left in the questionnaire and investigate a potential latent structure. After iterative item removal, the analysis led to a final structure of eSPS consisting of 12 items loading onto two correlated factors (r = 0.76, Figure 1). The first factor comprised eight items and closely resembled the human ‘Overstimulation’ subscale (e.g., ‘My horse tends to get emotionally overwhelmed’, My horse is unbothered when a lot is going on around him/her’, ‘My horse needs time to calm down after a stressful event’). The second factor comprised four items and was conceptually similar to the human ‘Sensitivity to Subtleties’ subscale (e.g., ‘My horse notices small changes in his/her everyday environment’ and ‘My horse reacts to everything he/she sees around him/her’).

**Figure 1.**
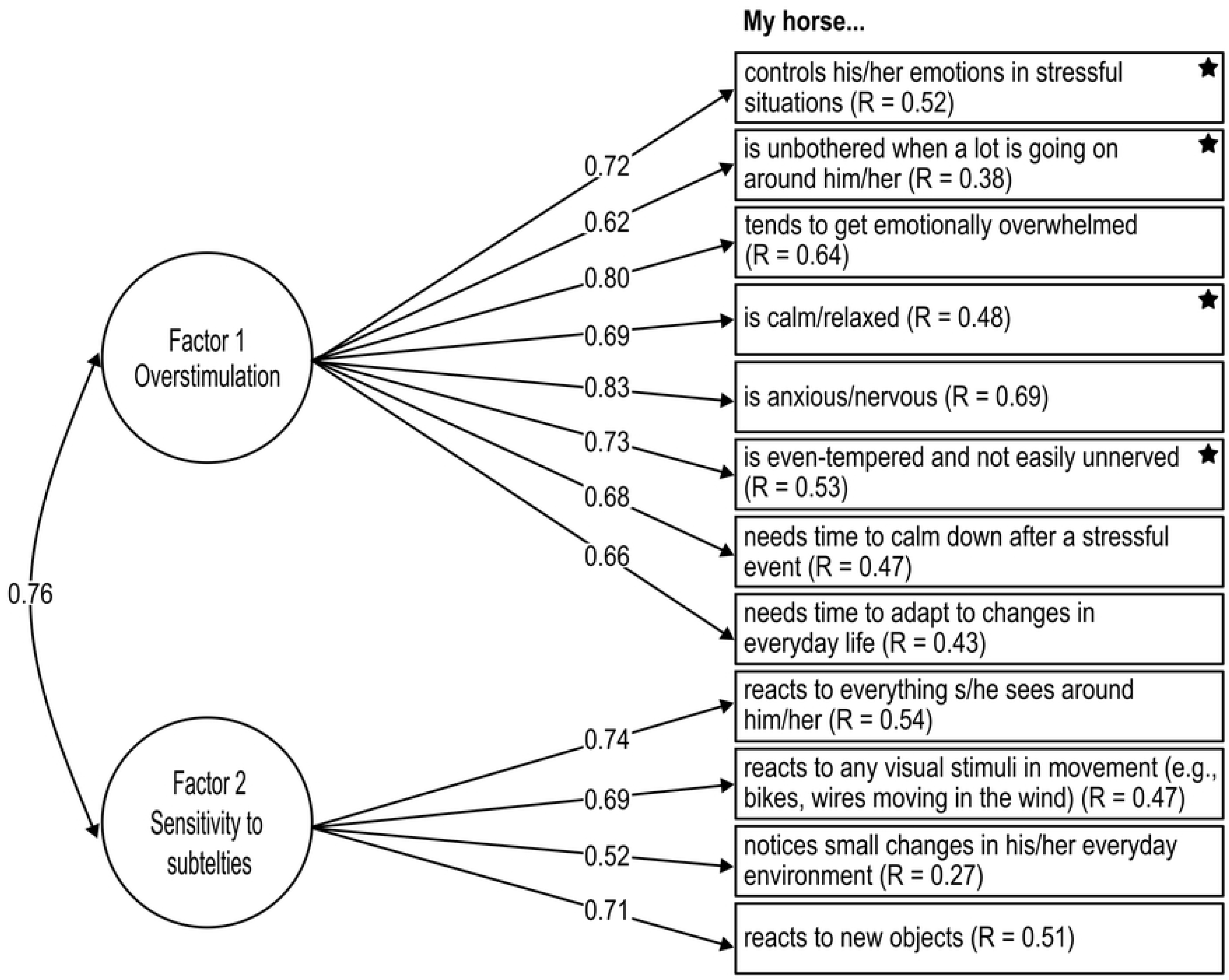
Path diagram showing the result from the Confirmatory Factor Analysis describing the structure of eSPS. Circles represent the two latent variables. The value on the two-headed arrow represents the correlation between them. The R value in each item represents how much of its variance is explained by the latent structure. Standardized factor loadings are presented along one-headed arrows and represent how items are predicted to vary if the latent variable changes. Stars represent items that were reverse coded so that a high value always indicate high levels of eSPS.

The final exploratory factor model showed acceptable to good fit (RMSR = 0.05, TLI = 0.91, RMSEA = 0.08, and off-diagonal fit index = 0.99). The two-factor structure was further evaluated using confirmatory factor analysis in the independent half of the sample and showed good fit (CFI = 0.96, RMSEA = 0.06, SRMR = 0.05).

On average, horses scored 3.30 on the first factor (SD = 1.26, range = 1.00–6.88) and 4.17 on the second factor (SD = 1.35, range = 1.00–7.00). On the whole scale, horses scored on average 3.59 (SD = 1.15, range = 1.00 – 6.83), with 25% of horses scoring above 4.33 (75^th^ percentile) and 10% of horses scoring above 5.17 (90^th^ percentile; Figure 2).

**Figure 2.**
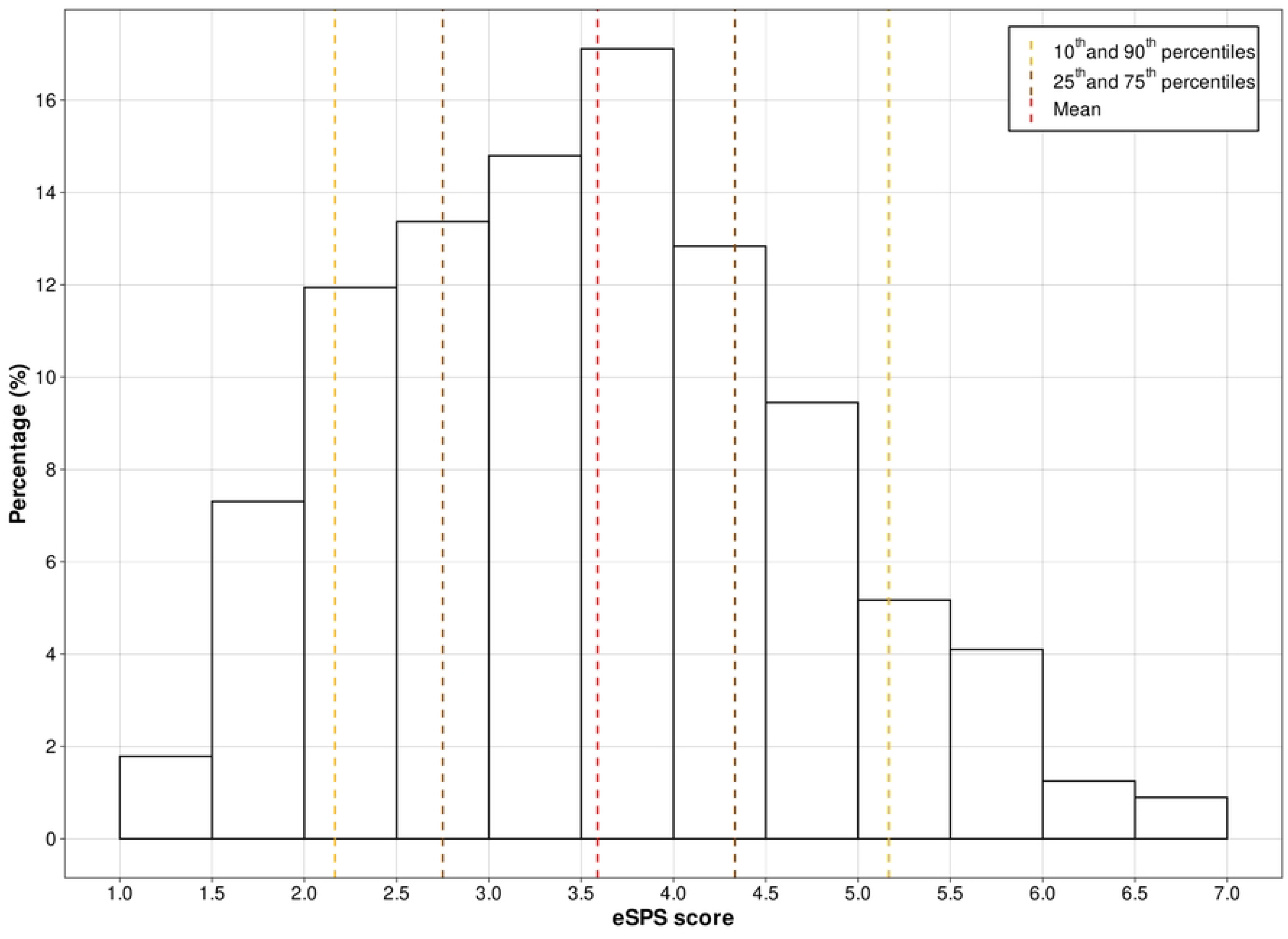
Distribution of eSPS score (calculated on the whole scale). The 10^th^ and 90^th^ percentiles are represented by dotted yellow lines. The 25^th^ and 75^th^ percentiles are represented by dotted brown lines. The mean is represented by a dotted red line.

Model comparisons indicated that the proposed two-factor model provided the best fit to the data. Specifically, the two-factor model yielded the lowest Akaike Information Criterion (AIC = 11’902), outperforming both the null model (AIC = 13’311) and the one-factor model (AIC = 11’980). Neither the second-order nor the bifactor model converged. In addition, internal consistency was high for the first factor, acceptable for the second factor, and high for the overall construct (Cronbach’s α = 0.89, 0.76 and 0.90, respectively). Inter-rater reliability was moderate, with an intraclass correlation coefficient of 0.67 (95% CI: 0.23–0.86), and intra-rater reliability was excellent, with an intraclass correlation coefficient of 0.90 (95% CI: 0.86–0.93).

The examination of convergent validity revealed that factor 1 of eSPS was strongly correlated with the personality traits ‘Apprehensive’ (r = 0.63), ‘Reliable’ (r = −0.65) and ‘Excitable’ (r = 0.74) described in [24] (Supplementary Table S3). Factor 1 also showed moderate correlations with ‘Fearfulness’ (r = 0.57), ‘Insecure’ (r = 0.56) and ‘Equable’ (r = −0.54). Factor 2 was only moderately associated with ‘Excitable’ (r = 0.51).

Results of linear models revealed that about 63% of the variance in eSPS was explained by five behavioural traits from [24] selected as broadly corresponding to dimensions commonly considered within the human Big Five framework, beyond age and gender, while 74% of the variance was explained when all behavioural traits from [24] were taken into account (Supplementary Table S4).

Incremental validity was assessed using owners’ responses to a demographic question indicating whether their horse exhibited behavioural problems. Adding the eSPS score to a model including 22 of the 25 behavioural traits described in [24] significantly improved the prediction of whether horses showed behavioural problem or not (ΔDeviance = 19.08, p < 0.001, Supplementary Table S5). The subsequent addition of the other three behavioural traits described in [24], which showed the strongest associations with eSPS (‘Apprehensive’, ‘Reliable’ and ‘Excitable’), resulted in a smaller but still significant improvement in model fit (ΔDeviance = 8.76, p = 0.033).

## Discussion

The present study aimed to investigate whether an equine Sensory Processing Sensitivity (eSPS) trait could be measured using an owner-reported questionnaire, and whether this construct represented more than a reformulation of previously described horse behavioural traits. Our results support both hypotheses. Through a systematic item selection and validation procedure, we identified a two-factor structure characterized by dimensions resembling the human SPS subscales of ‘Overstimulation’ and ‘Sensitivity to Subtleties’ [12]. To maintain consistency with the existing literature, we named our instrument the ‘Highly Sensitive Horse’ scale, following the naming conventions of previously developed scales for humans and dogs [8, 16]. The scale demonstrated satisfactory psychometric properties, including high internal consistency, excellent intra-rater reliability, and moderate inter-rater reliability. Although eSPS was associated with several established horse behavioural traits, it also explained unique variance in behaviour, suggesting that it captures aspects of individual variation not fully represented in existing equine personality frameworks. Our findings therefore provide empirical support that horses can be consistently evaluated in how they perceive and process environmental stimuli, extending previous behavioural evidence reported by [22].

The factor structure identified in this study shares notable similarities with SPS research in humans and dogs [9–13,16,17]. The first factor closely resembles the human and canine concept of overstimulation, describing individuals that become overwhelmed more easily and require more time to process challenging situations (e.g., ‘controls his/her emotions in stressful situations’, ‘needs time to calm down after a stressful event’) [12,13]. The second factor reflects heightened sensitivity to subtle environmental changes and increased attentiveness to surrounding stimuli (e.g., ‘notices small changes in his/her everyday environment’). Our results did not reproduce the full multidimensional structure proposed in human SPS models, particularly regarding emotional reactivity and depth of processing. However, this discrepancy should be interpreted cautiously, given that even in humans the dimensional structure of SPS remains actively debated [8,11,12,28,29].

Several explanations may account for the absence of factors corresponding to ‘emotional reactivity’ or ‘depth of processing’. These dimensions may truly be less differentiated in horses, or they may be more difficult for owners to observe reliably. While behavioural manifestations of overstimulation are often readily visible, cognitive processes such as ‘depth of processing’ are inherently more difficult to assess externally. Furthermore, owners’ capacity to perceive and categorize equine emotions and behaviours may vary with their level of experience, which could help explain why no distinct ‘emotional reactivity’ subscale emerged in our factor structure. While our questionnaire identified a sub-dimension related to ‘sensitivity to subtleties’, this factor contained fewer items than the overstimulation-related factor. This may reflect the difficulty for owners to assess their horses’ sensory experiences, such as what a horse smells, hears, sees, tastes, or feels.

These interpretations are further supported by the differences between the present scale and the canine SPS questionnaire [16]. The canine SPS scale contains substantially more items and captures a broader range of dimensions likely reflecting key differences in the human-animal relationship. Dog owners typically have the opportunity to observe their animals daily across a wide variety of contexts and become generally more familiar with canine behavioural signals, facial expressions, and body postures. Moreover, dogs have undergone a long history of co-evolution with humans, resulting in communication systems that are particularly accessible to human observers [30,31]. Horses, in contrast, have mainly been selected over their use and performance [32] and may therefore have experienced different selective pressures on human-directed communication. Horse owners also tend to spend less time observing their animals than dog owners do, further limiting access to the behavioural information required to evaluate subtle aspects of sensitivity. Differences in human–horse interactions and in the observability of equine behavioural signals may therefore contribute to the smaller number of dimensions captured by the present questionnaire. Assessing equine sensitivity through behavioural and cognitive testing may reveal additional dimensions of equine sensitivity that a questionnaire alone could not capture.

Overall, the psychometric properties of the scale provided encouraging support for its use. Internal consistency was high for both the overall scale and the two subscales, and intra-rater reliability was excellent. Inter-rater reliability was moderate, which was expected given that observers interact with the same horse in different contexts and possess varying levels of experience interpreting equine behaviour. Importantly, results suggest that eSPS should not be interpreted simply as another label for fearfulness or excitability. Existing behavioural traits explained only part of the variation in eSPS, and including eSPS in predictive models significantly improved behavioural predictions beyond previously established personality dimensions. The moderate correlations observed between eSPS and fearfulness and insecurity indicated that eSPS is at least partly independent from other behavioural traits, supporting its construct validity and being consistent with findings reported in humans and dogs [8,16]. The strong correlations with apprehensiveness, reliability, and excitability [24] were fully expected, as the definitions of these traits closely match the behavioural profile described by items loading on the first eSPS factor.

Several limitations should nevertheless be acknowledged. First, the study relied exclusively on owner-reported questionnaires, making it vulnerable to reporting biases and anthropomorphic interpretations [23]. Although considerable care was taken to minimize potential sources of questionnaire bias, some items may still have been open to interpretation. For example, the item ‘My horse needs time to calm down after a stressful event’ may be interpreted differently depending on the respondent’s experience and perception of equine behaviour. The moderate inter-rater reliability observed in our study suggests that ratings were reasonably consistent across observers, but that some variability in the interpretation of individual horses’ behaviour remained. This highlights an inherent limitation of owner-reported assessments and reinforces the need to complement questionnaire-based measures with standardized behavioural tests. Second, voluntary participation may have attracted respondents with a pre-existing interest in horse behaviour and personality, potentially limiting the representativeness of the sample. Third, the length of the online questionnaires may have contributed to participant fatigue and survey attrition (e.g., 24% of the people who started the questionnaire during the second online survey did not finish it).

An additional limitation concerns the initial item-reduction procedure. To reduce the number of items in the initial questionnaire, we classified horses as potentially highly or not highly sensitive using three screening questions and selected items that discriminated between these groups. Thus, the criterion used for initial item selection was not an independent or externally validated measure of eSPS and may have favoured items consistent with our initial conceptualisation of eSPS. However, this procedure was used only for preliminary item reduction. The retained items were subsequently evaluated in an independent and substantially larger online survey without using the initial sensitivity classification. The factor structure and validity analyses were therefore not influenced by the initial classification, reducing the risk that this procedure determined the observed structure of the final questionnaire.

Future research should now focus on behavioural validation of the eSPS construct. Questionnaire-based approaches provide valuable insights but remain constrained by the observer’s ability to perceive and interpret animal behaviour. Behavioural, cognitive, and physiological testing may reveal additional dimensions of sensitivity and facilitate comparisons across species. While questionnaires may be suitable for companion, farm or zoo animals, they are largely impossible to implement in most wild species. Developing standardized behavioural tests that assess individual differences in environmental sensitivity would provide definitive validation that SPS is a measurable, consistent, aspect of animal personality. Behavioural tests would further provide a more broadly applicable framework for investigating how different species perceive and respond to their surroundings, ultimately contributing to a comparative understanding of sensory processing sensitivity across the animal kingdom. Given the breadth of the analyses presented here, analyses of further validity were beyond the scope of this paper. However, we have begun the work of developing and performing objective behavioural testing and of investigating the links between eSPS and external and internal factors such as age, breed, sex, and detention modes.

## Conclusion

The present study provides evidence for a measurable equine Sensory Processing Sensitivity behavioural trait and introduces a validated questionnaire for its assessment. Although eSPS overlaps with existing behavioural traits, it also captures unique aspects of individual variation relating to how horses perceive and process environmental information. These findings suggest that understanding animal personality may require considering not only how individuals respond to the world, but also how they experience it. By incorporating variation in environmental sensitivity into personality research, future studies may gain a more complete understanding of individual differences in behaviour, welfare, and human-animal interactions.

## Methods

### Ethical statement

This study was reviewed and approved by the internal Research Ethics Committee of Agroscope (approval number: EK-AGS-2023-N04). All procedures involving humans were performed in accordance with the relevant ethical guidelines and regulations, including the ethical standards of the institutional research committee and of the 2024 Declaration of Helsinki. At each step, informed consent for participation and publication was obtained from all participants. For participants under 18 years of age, informed consent was obtained from a parent and/or legal guardian. Participants were provided with an information sheet outlining the purpose of the study, how data would be stored and used, the potential risks associated with their participation and their rights to withdraw. All responses were anonymized and stored securely on protected servers accessible only to the project team. Data will be retained for five years in accordance with the Swiss National Science Foundation policy and then securely destroyed. All data handling complied with the Swiss Federal Data Protection Act (FADP), and the European General Data Protection Regulation (GDPR). Licenses were obtained for the reproduction of Lloyd et al. [24] and Aron and Aron [8] questionnaires from Elsevier and the American Psychological Association, respectively. As the questionnaire is currently intended for research purposes, the authors consider it important to highlight that it should not be used yet as a decision-making tool for the selection, breeding, training or individual assessment of horses.

### Development of the questionnaire

To develop the Highly Sensitive Horse questionnaire, we used items from the validated Highly Sensitive Person, Highly Sensitive Child, and Highly Sensitive Dog questionnaires that could be express in a way appropriate for horses [8,9,16]. Since it was not possible to extrapolate all items, we additionally aimed at gathering a list of descriptors (i.e., short sentences describing personality aspects) of eSPS that are commonly used and understood by both professional and non-professional horse owners (following the approach from [16]).

To do so, we performed one-hour semi-directed interviews with horse owners with varying degrees of experience (N = 12 professionals and N = 8 non-professionals). We held the interviews in French, English and German, with a translator present when necessary, between the 27.11.2023 and the 30.01.2024. We interviewed owners based on a list provided by the Equidae Research Group of Agroscope, Swiss National Stud with owners who have had horses for a minimum of one year.

First, we asked owners to describe the behaviours of one or several of their horses they estimated to fit a highly sensitive profile: that is, horses who show a deeper processing of thought and emotions, greater likelihood of being over aroused, higher emotional intensity (both positive and negative) and higher sensitivity to subtle stimuli perceived by all modalities (visual, acoustic, tactile, olfactory, gustatory, proprioceptive). Expressions used by the owners to describe such horses included ‘tends to freeze and think for a long time before reacting to a situation’, ‘takes a long time to calm down’, ‘empathic’, ‘reacts vividly to touch’, ‘sees everything’.

Then, we specifically posed questions regarding the horses’ current living conditions, health, and behaviour (e.g., towards people and other horses, in stressful situations, when faced with novelty, etc.). We also encouraged owners to describe their horses’ personalities with their own words. Up until this point in the interviews, we ensured that we did not mention the words ‘sensitivity’, ‘sensory processing sensitivity’ or ‘hypersensitivity’ to avoid inducing a bias in the participants answers. The last two questions of the interview focused on whether the owners thought their horse might be highly sensitive, and, if so, what made them believe this to be the case, and whether they would consider the term ‘hypersensitivity’ suitable to their horse or whether other adjectives described their horse better. The aim of these questions was to ensure that the term was meaningful to all.

The combination of items extrapolated from the previously validated questionnaires and from the descriptors gathered during the interviews led to a questionnaire consisting of 104 items (see Supplementary Table S1 for all items and their sources). Again, we ensured that the words ‘sensitivity’, ‘sensory processing sensitivity’ or ‘hypersensitivity’ were not present in the questionnaire until the last item. We also followed guidelines from [23] to try to minimise potential biases (e.g., avoiding technical jargon, double-barrelled questions, insensitive measure). The initial questionnaire was developed in French. The items were then translated by two independent translators and back-translated by two other independent translators into both German and English in order to eliminate translation errors. All eight translators were native speakers.

### Refinement of the questionnaire

In a second step, we aimed at reducing the number of items in the Highly Sensitive Horse questionnaire to those that allow the distinction between different levels of sensitivity. This was done via an online survey. We published adverts on French, English, and German horse forums, as well as on the Facebook page of the Swiss National Stud asking horse owners to take part in ‘a horse personality questionnaire’. From this moment on, we defined ‘horse owners’ as the person who spends the most time with the horse whether it be the actual owner or e.g., a rider or handler. We published the Highly Sensitive Horse questionnaire on the online platform Tivian (unipark.com, Tivian XI GmbH, Cologne, Germany) between 24.05.2024 and 10.06.2024. We first ensured that owners knew their horse well enough to thoroughly answer the questionnaire by asking them to confirm that they had known their horses for at least six months and visited them at least twice a week. We then asked owners three questions to determine whether their horses might be on either of the extremes of the eSPS trait: 1. My horse perceives subtle external stimuli (e.g., when something in the environment slightly moves, makes a faint noise, lightly touches them, faintly smells, or has a subtle taste), 2. My horse picks up on my emotions/moods, 3. My horse is easily stressed. The questions were to be answered on a 3-point Likert scale to allow us to clearly separate horses into the highest *versus* lowest levels of sensitivity. Owners could then continue with the 104 items of the Highly Sensitive Horse questionnaire, which were to be answered on a 7-point Likert scale (1 = *strongly disagree*, 7 = *strongly agree*). ‘I don’t know’ and ‘Not applicable’ options were also added. Scores obtained from negatively worded items were inverted so that a higher score always indicated higher sensitivity (Supplementary Table S1). At this stage, the age of the participant was also recorded. Participants could only answer for one horse. In total, 121 owners returned fully filled questionnaires.

Using results from the three selective questions, we separated horses into two groups (‘potentially highly sensitive’ and ‘potentially not highly sensitive’) and statistically determined which items differed credibly between the groups (see ‘Statistical analysis’ below). These screening questions were used solely as preliminary criterion indicators for item reduction and were not included in the next validation step or in the final eSPS score. For the next step, we only kept items that showed a credible difference between the two groups (N = 45).

### Validation of the questionnaire

In order to validate the structure of the questionnaire, we performed a second large international online survey implemented on Tivian and available online between 02.07.2025 and 30.10.2025. We published adverts on the Facebook and Instagram pages of the Swiss National Stud asking horse owners to take part in ‘a horse personality questionnaire’. We also sent a similar solicitation to stables in France, Germany, the UK, and Switzerland whose email addresses were available online, as well as to horse owners who had previously participated in studies at the Swiss National Stud. Additionally, we posted adverts on Facebook group pages of horse owners in France, the UK and the USA.

The survey included 1) the Highly Sensitive Horse questionnaire (N = 45 items); 2) demographic questions such as: sex, age, country of residence, current living conditions of the horse, type of training methods used, health problems, behavioural problems, etc.; 3) previously validated questions about horse personality from [24]; and 4) the Highly Sensitive Person questionnaire from [8]. While the present study focuses on developing the Highly Sensitive Horse questionnaire, we added demographic questions and the Highly Sensitive Person questionnaire to the second online survey to allow future investigation of the relationships between equine SPS and external factors.

All but the demographic questions were presented in a random order to each participant and had to be answered on a 7-point Likert scale (1 = *strongly disagree*, 7 = *strongly agree*). Scores obtained from negatively worded items were inverted so that a high score always indicated high levels of the trait. We included the horse personality questionnaire here in order to distinguish eSPS from other behavioural traits such as fearfulness (following methodology from [8,16]). The survey was available in French, English and German. All items that had not already been translated in the previous stage were translated following the same methodology.

At the end of the survey, we asked participants whether they were willing to respond to the survey again six months later in order to assess intra-rater reliability. We also asked participants to name another person who closely knows their horse and invite them to complete the survey in order to assess inter-rater reliability. As opposed to the pilot questionnaire, participants in this survey had the opportunity to answer for as many horses as they wished. In total, 561 fully filled questionnaires were returned. After six months, 172 questionnaires were filled in again for intra-rater reliability, and an additional 23 persons took the questionnaire for inter-rater reliability evaluation.

### Statistical analysis

All statistical analyses were conducted in R [33] using the integrated development environment RStudio [34].

### 1. Refinement of the questionnaire

After the first step (‘Development of the questionnaire’), the questionnaire contained 104 items, which were administered online to horse owners. The objective was to identify which of these 104 items to retain in order to assess a horse’s level of eSPS. As an initial filtering step, we excluded items with more than 10% ‘I don’t know’ or ‘Not applicable’ responses, as this suggested that these items were not suitable or relevant for the broader equine population (N = 15).

Next, we classified horses into ‘potentially highly sensitive and ‘potentially not highly sensitive’ groups by first summing scores on the three initial screening questions designed to capture potential extremes of the eSPS spectrum. Based on the assumption that approximately 20% of humans exhibit high sensitivity and 30% exhibit low sensitivity [14], we classified the top 20% of horses with the highest scores as ‘potentially highly sensitive’ (N = 24) and the 30% of horses with the lowest scores as ‘potentially not highly sensitive’ (N = 36).

We fitted a Bayesian linear mixed effect model [35] following an ordinal distribution, with the score obtained on each of the 104 items as response variable. We added an interaction between the supposed group (highly *versus* not highly sensitive) and item number as fixed effect. This allowed us to investigate for each item if the score significantly differed between the groups. We also added owner’s language (French, English or German) and age as fixed effects, and horse ID as random effect. We used an uninformative prior and ran 65,000 iterations with a burn-in of 15,000, and selected every 50^th^ posterior parameter sample after the initial burn-in. We verified that the model assumptions were met by checking for the absence of autocorrelation (correlation between lags < 0.1; [35]), sufficient mixing (Markov-Chain-Monte-Carlo chain plots), and Heidelberg and Welch diagnostic test. To avoid prematurely excluding potentially informative items during this preliminary screening stage, we used a p-value threshold of 0.09 to identify items showing evidence of differences between the two groups, and thereby removed 43 items. We then calculated a correlation matrix between the remaining items. Only two items showed a high correlation (r > 0.8). We removed the item that had the highest occurrences of ‘Not applicable’ or ‘I don’t know’. The refinement of the questionnaire reduced the number of items from 104 to 45.

### 2. Validation of the questionnaire

We further validated the 45 selected items by administering them online again, together with demographic questions and previously validated questions about horse personality from [24]. Our aim was to examine whether the shared variance among items was driven by underlying latent variables, and whether the eSPS construct was at least partly independent from other equine behavioural traits.

First, we confirmed that no item showed near-zero variance. We then computed a Pearson correlation matrix to identify potentially problematic items, such as items showing correlations < 0.10 with all other items (which would therefore be unrelated to the overall construct) or items with correlations > 0.90 with at least one other item (which would be redundant). At this stage, no items were removed.

Second, we randomly split the dataset into two subsamples of comparable size (N = 280 and 281) so that we could perform exploratory factor analysis (EFA) on one half and confirmatory factor analysis (CFA) on the other half. Before we ran the EFA, we assessed sampling adequacy using the Kaiser–Meyer–Olkin (KMO) measure and Bartlett’s test of sphericity (KMO = 0.93; Bartlett’s test p < 0.05). We determined the number of factors using scree plot inspection [36] and parallel analysis [37], both of which supported a four-factor solution.

We conducted the EFA using minimum residual extraction and oblimin oblique rotation because we expected correlated factors. We treated the Likert-scale responses as continuous and used the raw data instead of a polychoric correlation matrix. Although responses remained ordinal, we followed common practice for seven-point Likert scales, since measurement distortion decreases when scales include five or more response categories [38]. We iteratively removed items based on three criteria: factor loading < 0.35, communality < 0.30, or complexity > 2.0. After each iteration, we ensured that each retained factor included at least three strongly loading items. When a factor failed to meet this criterion, we removed it. This procedure led us to a two-factor solution with 22 items. We then built a Pearson correlation matrix to detect multicollinearity patterns and to identify clusters of highly correlated items (r > 0.6). We removed problematic items and reran the EFA using the same procedure. This process produced a stable two-factor solution with a total of 12 items (Figure 1). We evaluated model fit using Root Mean Square Residual, Tucker-Lewis Index, Root Mean Square Error of Approximation (respectively RMSR, TLI, and RMSEA), and the off-diagonal fit index. To validate the factor structure, we ran a CFA on the second half of the dataset using the solution that the EFA had identified. We evaluated model fit using CFI, RMSEA, and SRMR (Standardized Root Mean Square Residual).

After we defined the structure of eSPS, we conducted additional validation analyses. First, we calculated internal consistency using Cronbach’s alpha for each factor and for the full construct. Second, we specified four additional CFA models (null, one-factor, second-order, and bifactor) using the same item set and compared their Akaike Information Criterion values to identify the best-fitting structure, which supported our two-factor solution.

Third, we computed scores by averaging items: a total eSPS score, a score for the first eSPS factor and a score for the second eSPS factor. We calculated Pearson correlations between these scores and established equine behavioural traits described by [24].

Fourth, we assessed incremental validity. To do so, we investigated whether the addition of eSPS score explained variance in the response to a demographic question asked in the questionnaire, namely whether the owner reported their horse to have a behavioural problem (e.g., biting, kicking, phobias) or not. We fitted four generalized linear models with a binomial distribution and the likelihood to show behavioural problem as response variable. Model 1 was a null model. Model 2 included behavioural traits from [24] which were not found to be strongly correlated with the eSPS score in the previous step. Model 3 added the eSPS score. Model 4 added the behavioural traits from [24] which were moderately or strongly correlated with the eSPS score. We compared models using ANOVA to test whether eSPS explained additional variance beyond established behavioural traits.

Fifth, we also examined how much variance in eSPS is explained by other behavioural traits using ANOVA. We fitted three linear models with eSPS score as the response variable. Model 1 included age and sex as predictors. Model 2 included age, sex, and five behavioural traits derived from [24]: Curious, Sociable, Excitable, Reliable, and Popular. These traits were selected based on a review of the horse personality literature [26,39–42] because they were considered to broadly correspond to dimensions commonly associated with the human Big Five personality framework. Although the personality structure proposed by [24] was not intended to map onto the human Big Five model, and indeed provides a more comprehensive description of equine personality, we included this comparison because previous research on sensitivity has frequently examined associations with Big Five personality dimensions. This approach therefore allowed us to place our findings within the broader sensitivity literature. Model 3 included all behavioural traits described by [24].

Finally, we assessed intra- and inter-rater reliability. For intra-rater reliability, we calculated mean eSPS scores per horse across two questionnaires completed by the same owner six months apart. For inter-rater reliability, we calculated mean eSPS scores per horse across owner and second-rater questionnaires. We computed intraclass correlation coefficients (ICC) using a two-way agreement model and interpreted them following [43], where ICC < 0.50 indicates poor reliability, 0.50–0.75 indicates moderate reliability, 0.75–0.90 indicates good reliability, and > 0.90 indicates excellent reliability. We also generated Bland–Altman plots and considered agreement acceptable when fewer than 5% of observations fell outside the limits of agreement (Supplementary Figure S6).

## Acknowledgments

The authors are grateful to all horse owners and operators who participated in the survey(s). The authors also thank the staff of The Swiss National Stud of Agroscope for their inputs regarding the content of the initial scale. This work was funded through a standalone grant (320030_213036) from the Swiss National Science Foundation awarded to SB.

## Supporting information

**Supplementary Table S1.** Initial Highly Sensitive Horse questionnaire and stages at which the items were removed. Items marked with an asterisk had their score reversed so that higher score always meant higher sensitivity. Items could be removed: because they had over 10% of “I don’t know” or “Not applicable” answer (stage 1a), because they did not help differentiate between potentially highly and non-highly sensitive horses (stage 1b), during the EFA process (stage 2 removal), or because they showed multicollinearity with other items (stage 1c or stage 2b). Items could originate from: interviews (I), Highly Sensitive Child (C), Highly Sensitive Person (P) or Highly Sensitive Dog (D) questionnaires.

**Supplementary Table S2.** Results from the Bayesian linear mixed effect model investigating the existence of significant differences between potentially “Highly Sensitive Horses” (HSH) and potentially “not Highly Sensitive Horses” (nHSH) for each of the 104 items of the Highly Sensitive Horse questionnaire. We present posterior mean, 95% confidence intervals (95%CI) and p-value. Results with p-value < 0.09 are written in bold. Results of the fixed items “language” and “age” together with results of the interaction between “item number” and “group” are presented here. Results of the fixed effects for each item and for each group are not presented for readability reasons but can be found at: https://osf.io/3dzr9/overview?view_only=68676324fc1940129b437d43ecb5cd04.

**Supplementary Table S3.** Pearson matrix correlation between personality items derived from Lloyd et al. (2007) and eSPS scores. Values above the grey cells are test value (r) and value below the grey cells are 95% confidence-intervals.

**Supplementary Table S4.** Anova results comparing linear models to assess how much variance in eSPS score is explained by other personality traits. The response variable for all models is eSPS score.

**Supplementary Table S5.** Anova results comparing generalized linear models to assess incremental validity. The response variable for all models is the likelihood for the horse to show behavioural problems (yes/no).

**Supplementary Figure S6.** Bland-Altman plots from inter-rater (A) and intra-rater (B) reliability estimation.

